# Who acquires infection from whom? Estimating herpesvirus transmission rates between wild rodent host groups

**DOI:** 10.1101/2020.09.18.302489

**Authors:** Diana Erazo, Amy B. Pedersen, Kayleigh Gallagher, Andy Fenton

## Abstract

To date, few studies of parasite epidemiology have investigated ‘who acquires infection from whom’ in wildlife populations. Nonetheless, identifying routes of disease transmission within a population, and determining the key groups of individuals that drive parasite transmission and maintenance, are fundamental to understanding disease dynamics. Gammaherpesviruses are a widespread group of DNA viruses that infect many vertebrate species, and murine gammaherpesviruses (i.e. MuHV-4) are a standard lab model for studying human herpesviruses, for which much about the pathology and immune response elicited to infection is well understood. However, despite this extensive research effort, primarily in the lab, the transmission route of murine gammaherpesviruses within their natural host populations is not well understood. Here, we aimed to understand wood mouse herpesvirus (WMHV) transmission, by fitting a series of population dynamic models to field data on wood mice naturally infected with WMHV and then estimating transmission parameters within and between demographic groups of the host population. Different models accounted for different combinations of host sex (male/female), age (subadult/adult) and transmission functions (density/frequency-dependent). We found that a density-dependent transmission model incorporating explicit sex groups fitted the data better than all other proposed models. Male-to-male transmission was the highest among all demographic groups, suggesting that male behaviour is a key factor driving WMHV transmission. Our models also suggest that transmission between sexes, although important, wasn’t symmetrical, with infected males playing a significant role in infecting naïve females but not vice versa. Overall this work shows the power of coupling population dynamic models with long-term field data to elucidate otherwise unobservable transmission processes in wild disease systems.

## Introduction

Understanding ‘who acquires infection from whom’ is a key challenge in epidemiology, population biology and disease ecology (1). For managing parasite species (defined here generally to include both macroparasites (i.e. helminths, ectoparasites) and microparasites/pathogens (i.e. viruses, bacteria, protozoans)) that can infect multiple host species, it is clearly important to establish both which host species play an important roles in transmission (2,3), and specifically why and how they contribute more to transmission than other species. Rabies transmission maintenance, for example, in the Serengeti ecosystem appears to be primarily dependent on domestic dogs, although it has the potential to infect all mammals, and its long-term persistence also typically depends on carnivores or bats (4). While it is clearly important to understand transmission between different groups of species for generalist parasites, it is also true for specialist parasites that infect only a single host species that it can be important to distinguish the contribution of different host population categories, such as age and sex classes, to determine which behaviours and groups are driving infection (5). For instance, a study of bank vole populations in north west England showed that cowpox virus-infected females might be a more important source of infection to either sex than are males (6). However, it can be notoriously difficult to identify routes of disease transmission and the key individuals or groups driving transmission in natural populations, which has limited our knowledge about who acquires infection from whom for many wildlife diseases.

Gammaherpesviruses are a widespread group of DNA viruses that infect vertebrates, including humans, and can lead to persistent infections (7). Gammaherpesviruses are largely host-specific (8), and as such murine gammaherpesviruses, such as murid herpesvirus 4 (MuHV-4, of which MHV-68 is the archetypal strain) have been commonly used in inbred laboratory mice (*Mus musculus*) as standard models for understanding gammaherpesvirus infection biology more generally (9,10). However, while the mechanisms of pathogenesis and immune interactions within individual hosts have been well studied (10,11), for instance it is well known that infection fluctuates between acute and latent infection stages, but only acute individuals transmit the virus (12), little is known about the routes of transmission in either the laboratory setting or in natural host populations.

It is widely considered that gammaherpesviruses are transmitted through close contact (13). Nonetheless, previous laboratory studies in female mice infected with murid gammaherpesvirus-4 (MuHV-4) through an intranasal inoculation failed to produce any new infections of naïve mice in the same cage, suggesting that transmission does not happen via general contact followed by inhalation of airborne virus particles (14). Furthermore, male presence and natural behaviour could be critical in virus transmission(15), leading to the hypothesis that infection could occur through sexual contact and or aggressive encounters. Empirical studies in wild populations have shown that older males were more likely to be seropositive for gammaherpesviruses compared to other groups within the population, with the highest rates of seropositivity in males with high body mass (15,16). Knowles *et al*. (2012) highlighted the importance of reproductive status, suggesting that reproductively active males are more likely to be infected than non-reproductive males, whereas females showed no evidence for any effect of reproductive status. The evidence thus far suggests that behaviours specific to reproductively active males could be driving herpesvirus transmission, supporting the hypothesis of transmission maintained by male dominance behaviours such as biting and scent marking behaviour. However, one laboratory experiment successfully demonstrated transmission of murine gammaherpesvirus-68 (MHV-68) from infected females to naive males after sexual contact (17), so the primary directionality of transmission between males and females remains unclear.

The route of transmission, and the behaviour involved, have important implications for how transmission likely scales with population size. If strict sexual transmission is the main driver of murine gammaherpesviruses infection, we would expect transmission to be mainly frequency-dependent, because individuals generally do not have a greater number of sexual contacts in higher-density populations (18,19). However, if transmission is primarily driven by male dominance behaviours (biting and scent-marking) then transmission may be primarily density-dependent. However, there is debate on whether density-dependent transmission depends on the overall population density or not. A previous study of WMHV in wild wood mice did not find strong density-dependent patterns, because prevalence was not significantly associated with an increase in the total population during the breeding season (16). Another study revealed that population density increased from July onwards, but WMHV prevalence remained low (15). Thus, if male dominance behaviour is maintaining herpesvirus infection in wild rodent populations it is likely that transmission: i) depends mostly on male density, particularly reproductive males, and ii) occurs before breeding season when reproductively active males are establishing territories, and the number of contacts with other reproductive males increases. Overall then, although previous studies (15,16) suggest an asymmetrical role of males in herpesvirus transmission, uncertainty remains about what are the key disease transmission routes and the contribution of each population category.

To elucidate key aspects of transmission biology which would be otherwise hard to identify, mechanistic model fitting to data can be a useful tool that aids detection and quantification of potential transmission routes (20). Here, we investigate WMHVs transmission dynamics within its natural host population by fitting a series of mechanistic models to 4-years of longitudinal serological data from a wild population of the wood mouse, *Apodemus sylvaticus*. Each model reflects the above-mentioned hypotheses about the transmission processes within and between demographic groups by incorporating different combinations of host sex (male/female), age (subadult/adult) and transmission mechanisms (density/frequency-dependent). We compare models based on the best fit and analyse the estimated parameters, including the different group contributions to the overall reproductive number (R_0_) of the virus. Finally, we discuss the implications of our results in the context of WMHV infection, aiming to shed light on its natural transmission mechanism.

## Methods

### Field data

Serological data from longitudinal, repeat sampling of wood mice (*Apodemus sylvaticus*) collected between June 2009 and December 2012 in four grids located in Haddon Wood, Cheshire, UK as described in (15). Briefly, two Sherman live-traps were baited with grain and bedding and placed every 10 m in each 70 x 70 m squared grid. Trapping was conducted for 3 consecutive nights every 3 weeks during four annual field seasons: June-December in 2009 and 2012, and May-December in 2010 and 2011. All trapped individuals were tagged with a unique passive integrated transponder (PIT) tag for recognition in subsequent recaptures. From all wood mice, morphometric measures were recorded, including sex and age (juvenile, sub-adults, and adult), and blood samples were taken from the tail at first capture within each month for serological analyses. We used a serological assay that detects WMHV antibodies in mouse serum using IFA (15). Since maternal antibodies to WMHV persist into young adulthood, we excluded juveniles from our analysis.

### Models

We fit a series of mathematical models to the serological data, each capturing different potential aspects of WMHV transmission (see below). All models used the same demographic framework to describe the wood mouse population dynamics, against which WHMV transmission occurred (i.e., mouse population dynamics were assumed to be independent of viral infection dynamics, implying a negligible impact of infection on host survival or reproduction). Hence, we first fitted a demographic model to the observed mouse population dynamic data, and in doing so estimated the key demographic parameters (birth rates, survival rates and carrying capacities) for our specific system. For this demographic component, we constructed a two-equation model representing wood mice population dynamics, as follows:

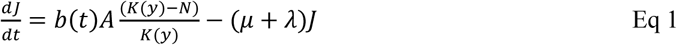

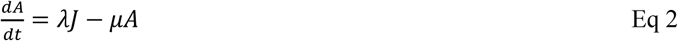

where *J* represents the juvenile and sub-adult mice combined, *A* the adult mice, *N* the total mice population size, *K*(*y*) the carrying capacity in year *y, b*(*t*) the birth rate, *μ* the mortality rate and *λ* the maturation rate. Wood mice population dynamics are highly seasonal in UK woodlands, thus we defined birth rate as a function of time, as follows:

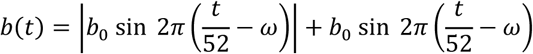

where *t* is time in weeks, *b*_2_ is the baseline birth rate, *ω* is the birth function phase (which sets the timing of peak births). This construction means *b*(*t*) is periodic with a period of 1 year, allowing for a 6-month-long breeding season and 6 months outside that period of no reproduction which fits patterns of breeding in the field sites. The carrying capacity, *K*(*y*), was also assumed to vary for each of the four years of our study, to account for year-to-year fluctuations in overall mouse population size, dependent on seasonal environmental factors (e.g. resources or abiotic conditions); hence we assumed different values of *K*(*y*) for each year, which we estimated through model fitting (see below).

Having estimated the relevant parameters to describe the wood mouse population dynamics, we then used that demographic model as a baseline for the construction of four herpesvirus epidemiological models, which we fitted to the observed serological data, estimating the key epidemiological parameters in the process. These models accounted for previous findings that suggest WMHV and the related MuHV-4 infection are sex- and age-differentiated (15,16), and hence different age and sex classes may contribute differently to transmission. The basic herpesvirus model (Figure 1A) consisted of a three-equation system with susceptible, active, and latent classes, and just one age class (i.e., the adult/sub-adult distinction from Eq1 is subsumed into a single equation of overall host dynamics):

**Figure 1.**
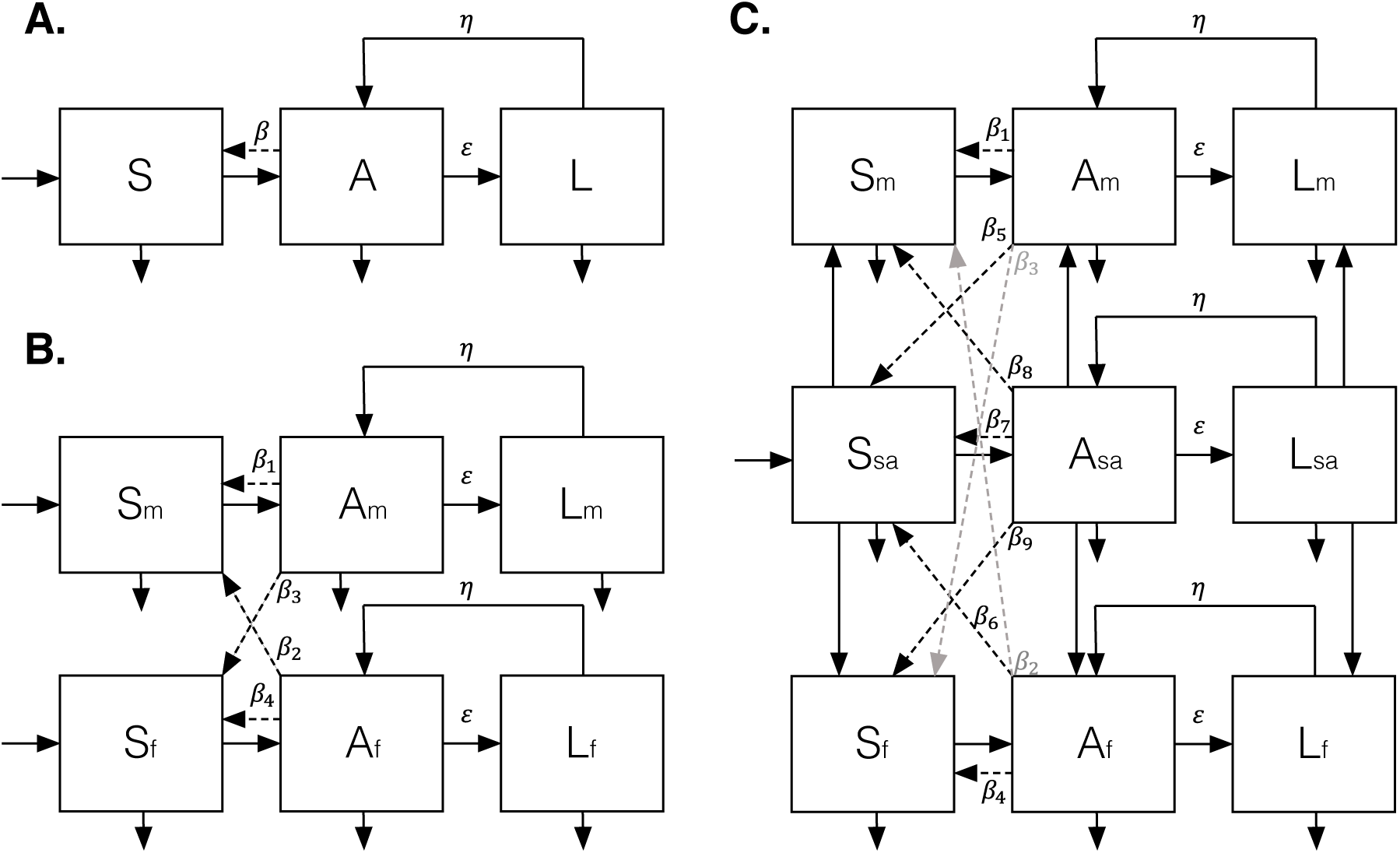
Herpesvirus models. A) Basic model. B) Sex-explicit models: six group density-dependent model (*6-group DD*) and six group frequency-dependent model (*6-group FD*). C) Sex- and age-explicit models: nine group density-dependent model (*9-group DD*) and nine group frequency-dependent model (*9-group FD*).

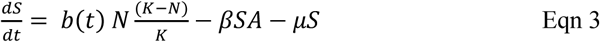

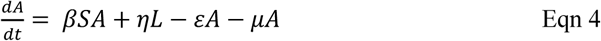

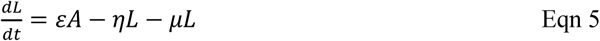

Susceptible individuals (*S*) become infected and enter the active infection class (*A*) through contact with active-infected individuals (latent-infected individuals are assumed not to transmit infections) at transmission rate *β*. Active-infected individuals then move to the latent class (*L*) at transition rate *ε* (hence active infections last on average 1/*ε* weeks), and vice versa at rate *η* (hence latent infections last on average 1/*η* weeks). Based on existing knowledge about gammaherpes viruses (8) it is assumed individuals do not recover from infections. All individuals, whether uninfected, active infected or latent-infected, were assumed to die at the same rate *μ*.

Using the above framework as a baseline, we developed a series of alternative formulations that incorporated potential transmission pathways for each specific demographic group in the population, and different formulations of the transmission functions. The first model (six demographic groups with density-dependent transmission: *6-group DD*) considered explicit female and male classes (Figure 1B) and density-dependent transmission for all possible transmission routes (*β*_1_ − *β*_4_). Therefore, the model consisted of six groups (susceptible, active-infected and latent-infected, for both females and males), and all transmission terms had the form *β*_*n*_*S*_*i*_*A*_*j*_ where *i* and *j* represent sex, either female or male, and *β*_*n*_ is the transmission rate between *i* and *j*. The second model (six group frequency-dependent transmission: *6-group FD*) had the same structure as the *6-group DD* model but considered frequency-dependent transmission (as is often assumed for sexually transmitted infections; (19)), thus the transmission terms had the form 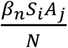. The third (nine group density-dependent transmission: *9-group DD*) and fourth (nine group frequency-dependent transmission: *9-group DD*) models accounted for age-dependent transmission by explicitly incorporating a subadult class into the above framework, along with female and male adult classes (Figure 1C). Based on our previous results (15), we assumed that male and female subadults do not differ in terms of transmission potential or susceptibility, and so were combined into a single class. Subadults were assumed to mature to adults at rate *λ* (see demographic model, above), half becoming adult males and half adult females. This structure therefore introduced five additional possible transmission routes involving the subadults, resulting in nine transmission parameters (*β*_1_ − *β*_9_) overall. As with *6-group DD* and *6-group FD* models, *9-group DD* and *9-group FD* models differed by their transmission function; *9-group DD* used density-dependent transmission for all transmission rates (*β*_*n*_*S*_*i*_*A*_*j*_) and *9-group FD* used frequency-dependent transmission for all transmission rates 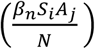. See Table 1 for a description of all parameters in all models.

**Table 1A.** Parameters and values used in the demographic model. Estimated values show median and 95% credible intervals from the posterior distributions of each parameter.

| Symbol | Parameter description | Baseline value | Prior distribution | Estimated value |
| --- | --- | --- | --- | --- |
| $b_0$ | Birth function scale | 0.5 | 0.5 – 20 | 0.539<br>[0.538 – 0.540] |
| $\omega$ | Birth function phase | 0.15 | 0 – 1 | 0.0148<br>[0.0145 – 0.0151] |
| $\mu$ | Mice mortality rate | 0.04 | 0 – 1 | 0.0596<br>[0.0595 – 0.0597] |
| $\lambda$ | Maturation rate | 0.1 | 0 – 1 | 0.102<br>[0.101 – 0.103] |
| $K(1)$ | Carrying capacity in year 1 | 50 | 25 – 100 | 81.3<br>[81.2 – 81.4] |
| $K(2)$ | Carrying capacity in year 2 | 50 | 25 – 100 | 56.8<br>[56.7 – 56.9] |
| $K(3)$ | Carrying capacity in year 3 | 50 | 25 – 100 | 95.5<br>[95.4 – 95.6] |
| $K(4)$ | Carrying capacity in year 4 | 50 | 25 – 100 | 69.2<br>[69.1 – 69.3] |

### Model parameterization, fitting and comparison

Demographic parameters (θ_1_ = {*K, μ, λ, b*_0_, *ω*}) and disease-related parameters (θ_2_ = {*β*_1_ … *β*_9_, *ε, η*}) were estimated using adaptive Monte Carlo Markov Chain Metropolis-Hastings (MCMC-MH), assuming uniform priors (Table 1A), through fitting to data on mouse population abundance and infection seroprevalence respectively. We ignored the first year (52 weeks) of predicted transient dynamics of the simulation as burn-in time and fit the models over the subsequent 4 years of data.

As described above, model fitting was carried out in two stages. First, demographic parameters and yearly carrying capacities were estimated by fitting the simulated total number of weekly mice (*l*_*mice*_; *J* + *A* from Eqn 1 and 2) to the observed number of mice captured per week (*y*_*week*_). The weekly number of mice captured was assumed to follow a Poisson distribution. The log-likelihood of the data for the mice population dynamics model was given by:

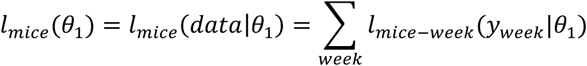

Through this we generated posterior distributions for the 4 year-specific carrying capacities (2009-2012), baseline birth rate and birth function phase.

Next, we sought to estimate the different transmission parameters in the various models. For model fitting to data the simulated WMHV prevalence of wood mice in each demographic group (proportion of animals infected per group each week) was contrasted to the observed prevalence in each group per week; since the data on infection status were based on a serological assay, which cannot differentiate active from latent infections, we combined the predicted prevalences in the *A* and *L* classes (Eqns 4 and 5) to calculate an overall expected prevalence of infection in each group (e.g., the proportion of predicted infected males was given as (*A*_*male*_ + *L*_*male*_)/(*S*_*male*_ + *A*_*male*_ + *L*_*male*_)). Observed weekly prevalence was assumed to follow a binomial distribution. The overall log-likelihood of the data for the first and second model (only females and males) was given by:

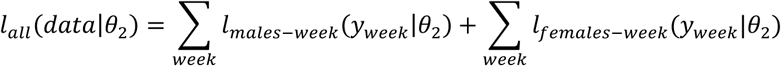

The overall log-likelihood of the data for the third and fourth model was given by:

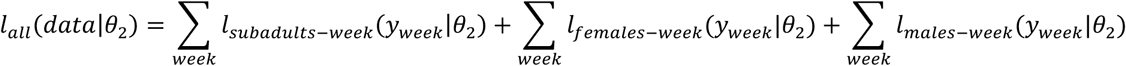

Through this we generated posterior distributions for disease-related parameters (the relevant transmission rates, *β*_*i*_, and the transition rates, *ε* and *η*).

For each model, we ran four MCMC-MH chains of 10,000 iterations using the default parameter standard deviation (parameter value divided by 10). Then, for each chain, using the first chain output (standard deviation and 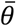 for the last 9,000 iterations) as input, we ran a second chain of 100,000 iterations. For the second chain, the first 5,000 iterations were discarded (burn-in), and we eliminated every 10 samples per sample to avoid auto-correlation (thin). The Gelman-Rubin diagnostic was used to assess MCMC convergence by analysing the difference between chains for each model. Thinned and burned chains were combined for comparison between the herpesvirus models.

Herpesvirus models (*6-group DD, 6-group FD, 9-group DD* and *9-group FD*) were compared using the total Deviance Information Criterion (DIC) for each model; the model with the lowest DIC values corresponds to the best model. Having estimated the epidemiological parameters for the best-fitting model, we used those values to derive an estimate of the overall basic reproductive number, *R*_*0,tot*_, using the next generation method of Diekmann, *et al*. (2010) (21), and partitioned that value into the contributions of males and females, respectively.

All model fitting and comparisons were carried out using R version 3.5.1 and the RStudio Integrated Development Environment (IDE) (22). Adaptive Monte Carlo Markov Chain Metropolis-Hastings (MCMC-MH) was conducted using the fitR package (23) and outputs were analysed using the coda package (24).

## Results

Overall during the four-year field sampling, 1,394 mice were captured a total of 4,316 times, with a mean number of captures per mouse of 3.04 (range 1− 28). A total of 1,343 mice were identified as subadults or adults and were captured 4,197 times. The remaining 51 mice were identified as juveniles. The number of subadult and adult mice considered in this study were 1,065, for a total of 1,817 samples. Captures with missing data or that did not have blood sample taken, were not considered. Overall, 15.4% animals were seropositive for WMHV, with a significant difference in WMHV incidence between the sexes (0.1 of female captures were seropositive, compared to 0.19 of male captures; *X*^2^ = 19.79, *p* < 0.001).

Regarding the estimation of demographic parameters, the birth function scale (*b*_0_) and phase (*ω*) were estimated to be 0.539 and 0.0148, respectively, implying on average 17.8 offspring per individual per breeding season and that the population is expected to increase at the beginning of the field season, corresponding well to the observed trends in the data (Figure 2). We estimated that mean wood mouse life expectancy was approximately17 weeks (*μ* = 0.0596 [0.0595 − 0.0597] week^−1^) and mean maturation time to sexually reproductive status was around 10 weeks (*λ* = 0.102 [0.101 − 0.103] week^−1^). The highest estimated carrying capacity corresponded to the third field season (2011) with 95 mice, followed by the first and fourth field seasons (2009 and 2012) with 81 and 69 mice, respectively. The lowest carrying capacity was estimated for the second field season (2010) with 57 mice. See Table 1A for 95% credible intervals from the posterior distributions of demographic parameters.

**Figure 2.**
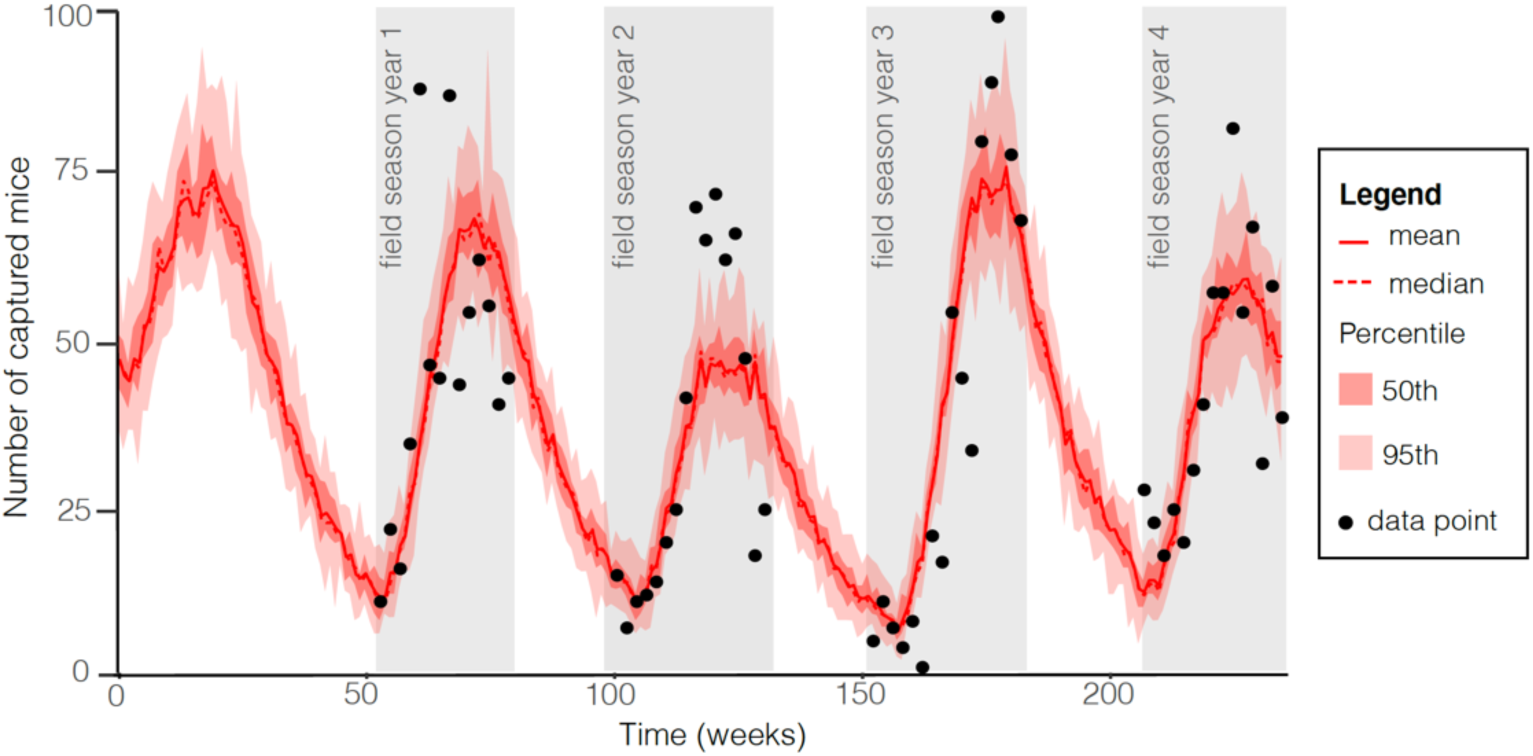
Demographic model fit to data. *X-*axis is time in weeks from the start of the simulation (the first 52 weeks were excluded in model fitting as burn-in time), and *y-*axis is number of captured mice. Dots represent data points and grey shaded regions cover the period in which fieldwork was conducted. Model simulation mean and median are shown by the red solid and dashed lines, respectively. Pink and red shades represent 50^th^ and 95^th^ percentile of the simulations.

The best-fitting transmission model by DIC (Table 2) incorporated both female and male classes explicitly, but not subadults, and assumed density-dependent transmission (*6-group DD* model). These results suggest that the most accurate and parsimonious description of the observed patterns of WMHV transmission in these wood mouse populations requires explicit description of transmission within and between the two sexes, but accounting for age-varying transmission processes was not necessary. This best-fitting model showed good agreement with the observed weekly prevalence for both adult females and males (Figure 3), capturing the observed infection peaks during the four seasons, for females and males during June 16 of 2009 (week 53), June 1 – June 29 of 2010 (weeks 103 – 107), June 14 of 2011 (only males, week 157), and July 3 – 31 (weeks 212 – 216). Interestingly, the third field season (2011) showed the lowest infection peak and the highest carrying capacity among seasons, and the second season had the lowest estimated carrying capacity. An increase in prevalence was observed between December to May-June for males and females in the data and the model simulation (Figure 3), except females from year 2 to year 3 (2010 – 2011), because at the beginning of year 3, the number of infected females was zero.

**Table 2.** DIC values comparing model fits for each demographic group, and overall, for the four fitted models.

|  | Female | Male | Sub-adults | Total |
| --- | --- | --- | --- | --- |
| **Model 1: six group density-dependent transmission** | 179 | 298 | - | 463 |
| **Model 2: six group frequency-dependent transmission** | 491 | 609 | - | 792 |
| **Model 3: nine group density-dependent transmission** | 173 | 253 | 91 | 492 |
| **Model 4: nine group frequency-dependent transmission** | 409 | 519 | 320 | 774 |

**Figure 3.**
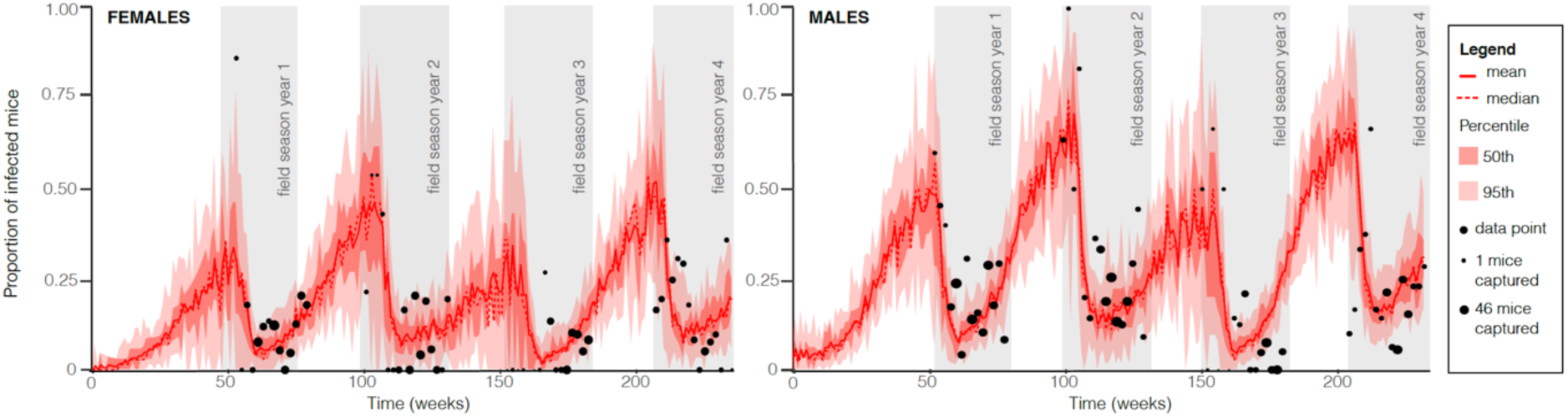
Six group density-dependent model (*6-group DD*) fit to data. Left and right panels show model fit to female and male data, respectively. *X-*axis is time in weeks from the start of the simulation (first 52 weeks were excluded in model fitting as burn-in time) and *y-*axis is herpesvirus seroprevalence (proportion seropositive; for the simulation these comprised both acute and latent infections). Dots represent data points and dot size illustrates the number of mice captured at each data point. Areas in grey shade show the period in which fieldwork took place. Model simulation mean and median are shown by the red solid and dashed lines, respectively. Pink and red shades represent 50^th^ and 95^th^ percentile of the simulations.

Among the estimated transmission rates for the best-fitting model (*β*_1_ − *β*_4_), transmission from males dominated; male-to-male transmission was the highest (*β*_1_ = 0.00789 [0.00786 − 0.00791]; Table 1B), followed by male to female transmission (*β*_3_ = 0.00322 [0.00321 − 0.00323]). Conversely, transmission from females, either to males (*β*_2_ = 0.00091 [0.00090 − 0.00092]) or to other females (*β*_4_ = 0.00167 [0.00165 − 0.00169]), was very low and on average 77% lower compared to transmission from males. The estimated transition from active to latent infection was rapid (*η* = 0.624 [0.627 − 0.621] per week), implying a median duration of the initial active infection phase of ∼1.6 weeks. The estimated median transition time back from latent to active infection was ∼2.5 weeks (*ε* = 0.403 [0.4 − 0.406] per week). Parameter estimates for the other herpesvirus models can be found in Table 1B.

**Table 1B.**
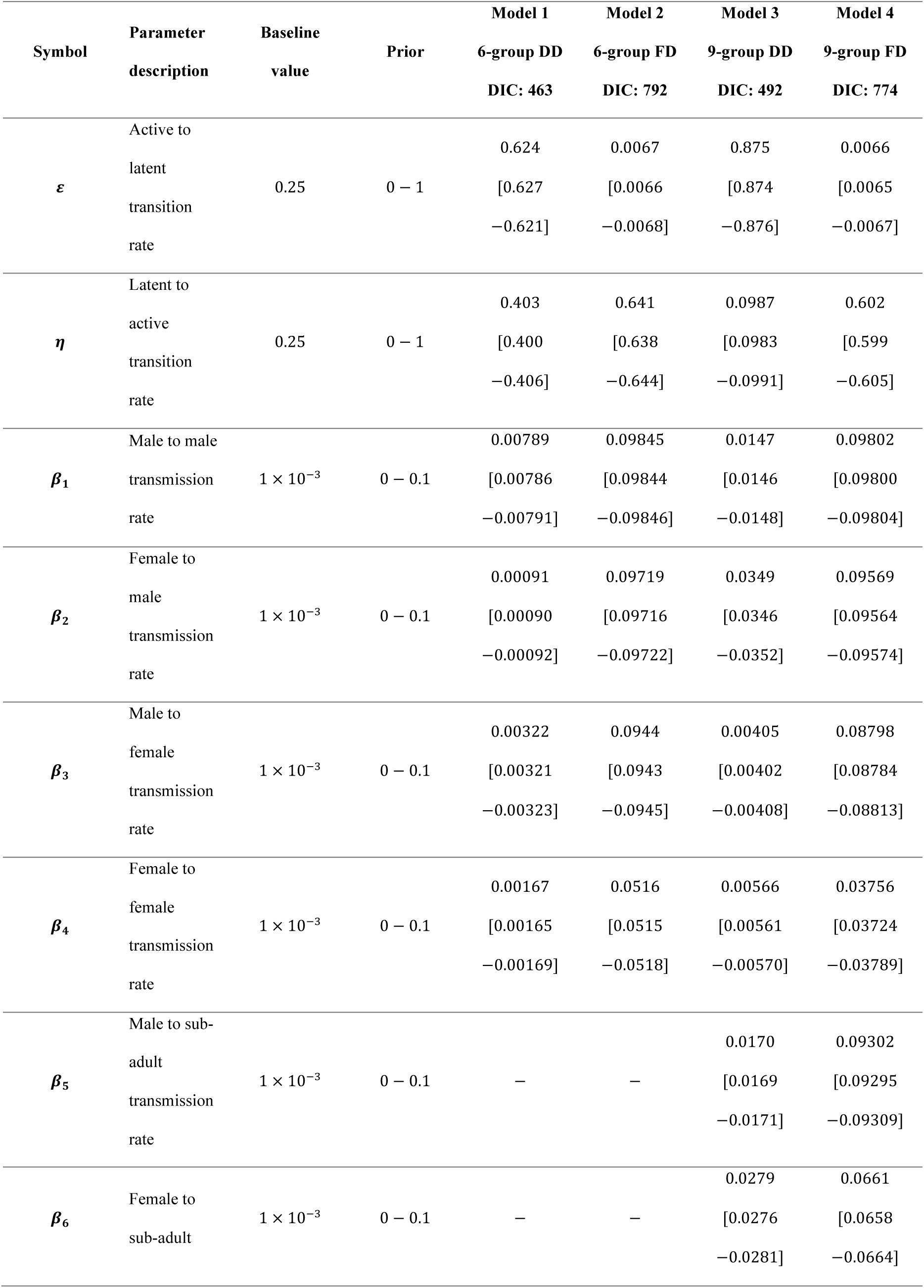
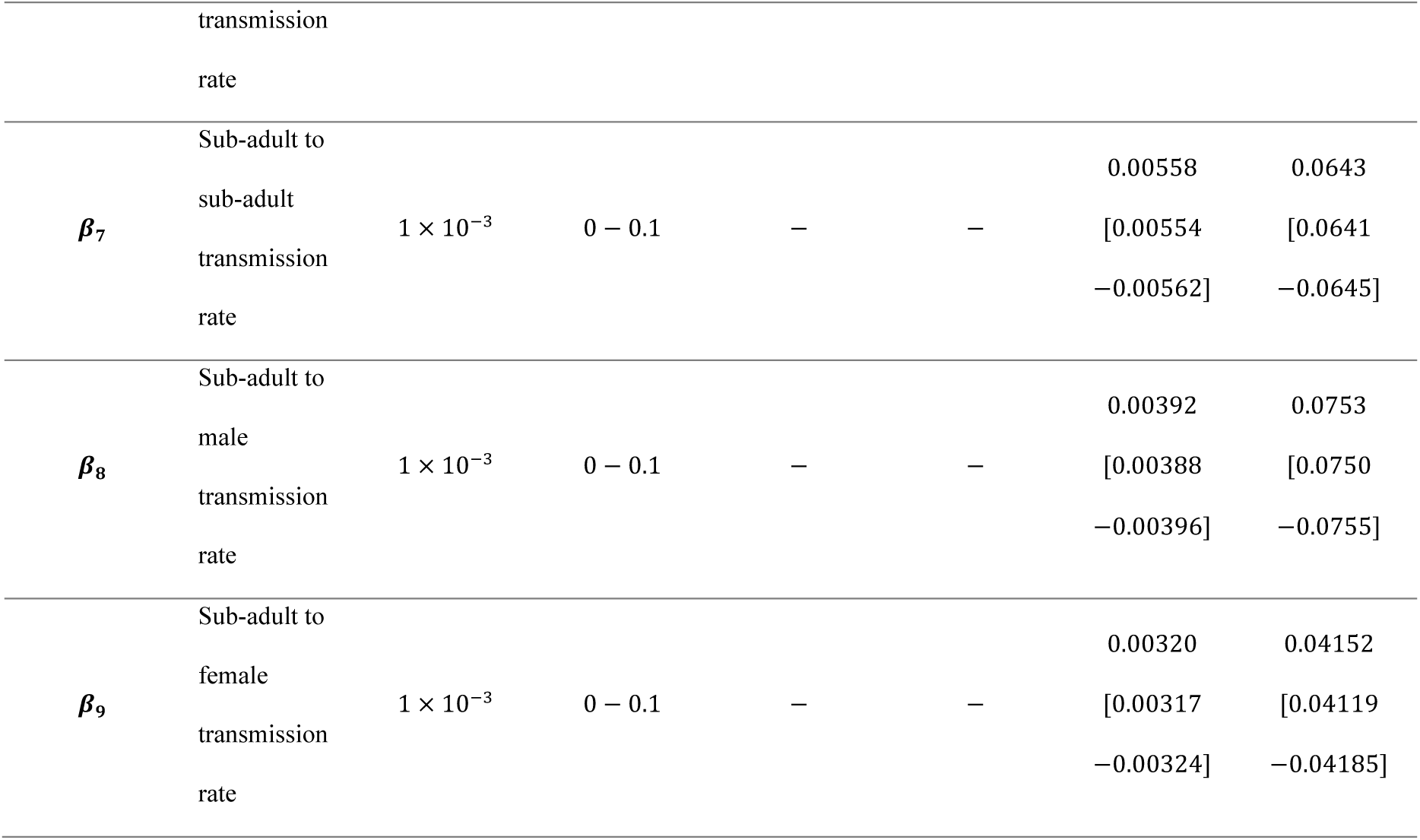
Parameters and values used in the four herpesvirus models: *6-group DD* and *6-group FD, 9-group DD* and *9-group FD*. Values show median and 95% credible intervals from the posterior distributions of each parameter.

Using the *6-group DD* model estimated values we assessed the contribution of males and females to the overall basic reproductive number (*R*_*0,tot*_) for WMHV. Following Diekmann, *et al*. (2010), we calculated an expression for the *6-group DD* model *R*_*0,tot*_ using the next generation matrix (NGM) approach (21), where *R*_*0,tot*_ is represented the dominant eigenvalue of a matrix resultant from the multiplication of the transmission and the transition matrix. The transmission matrix *T* represented the production of new infections as follows:

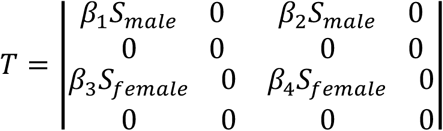

and the transition matrix Σ, describing changes in state (including death), was given by:

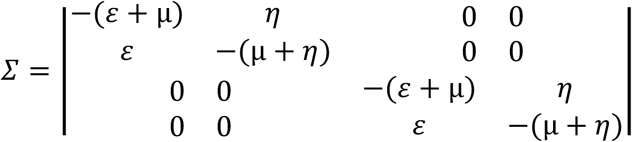

*R*_*0,tot*_ is then dominant eigenvalue of the next generation matrix approach given by −*T*Σ^−1^:

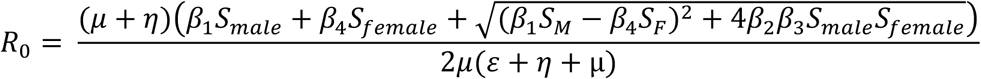

Assuming a 1:1 female:male ratio, the total number of mice as the mean carrying capacity over the four-year period 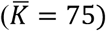, and replacing with the estimated values from the best-fitting *6-group DD* model (Table 1B), we estimated *R*_*0,tot*_ as 2.1299 [2.1290, 2.1307].

Finally, we partitioned *R*_*0,tot*_ into the contributions of males and females by setting either the male-onward (*β*_1_ and *β*_2_) or female-onward (*β*_3_ and *β*_4_) transmission rates to 0. In other words, this procedure calculated R_0_ when either males or females contributed nothing to transmission, resulting in: *R*_0,*males*_ = 2.0133 [2.011 − 2.0153] and *R*_0,*females*_ = 0.4222 [0.4186 − 0.4276]. Note that, since there are individuals that could get infected by both groups, leading to potential for overlaps in transmission, *R*_0,*males*_ and *R*_0,*females*_ do not simply add up to *R*_*0,tot*_. Following the conceptual framework of Fenton *et al*. (2015) (3), and using these *R*_0,*males*_ and *R*_0,*females*_ values, Figure 4 shows that males can maintain WMHV on their own, and infections in females are primarily due to spillover from males to females.

**Figure 4.**
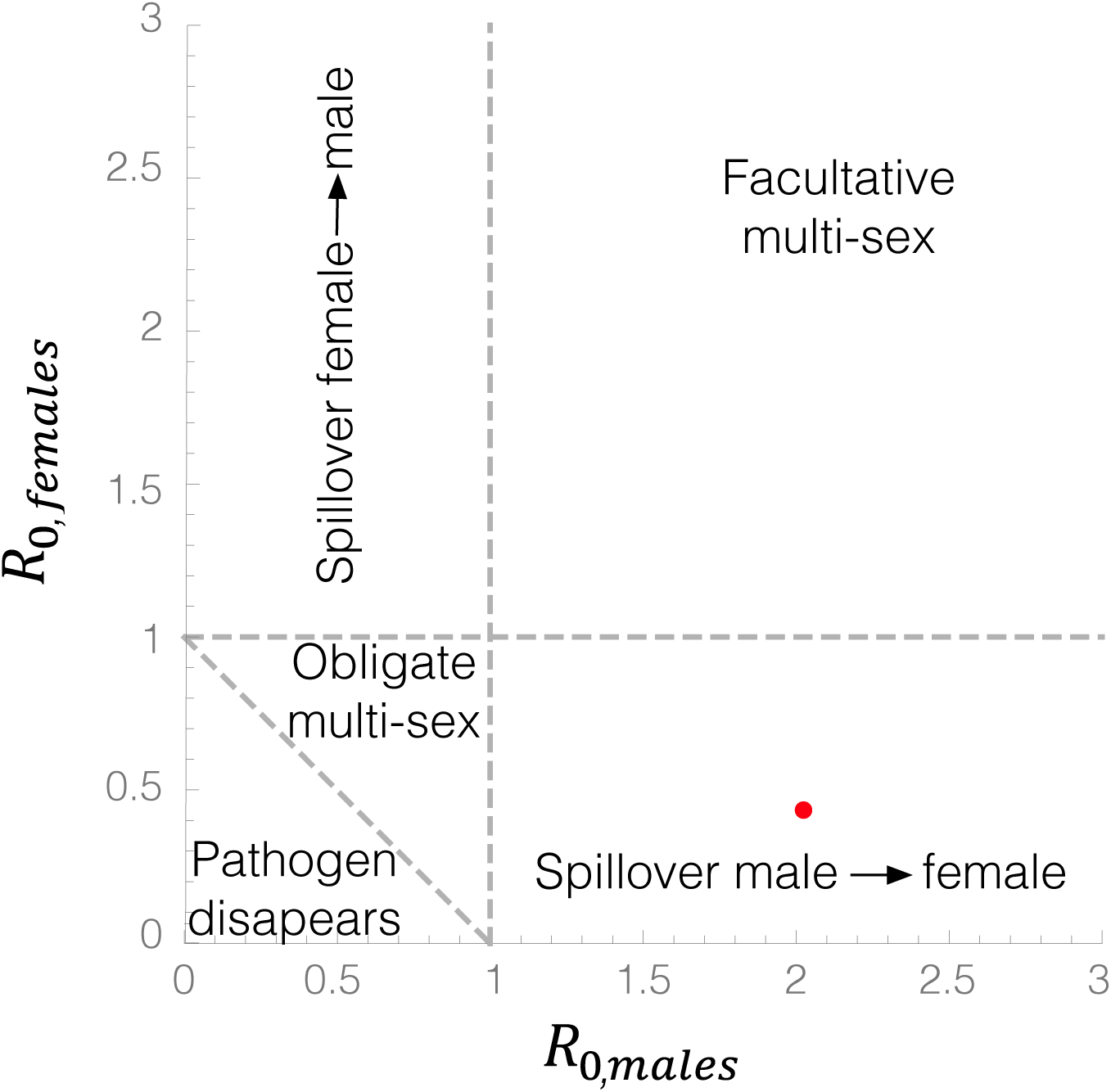
*R*_0,*males*_ - *R*_0,*females*_ parameter space. Based on the framework by Fenton et al (2015) (3), the figure has five regions, each representing a potential disease outcome: parasite exclusion, spillover from male to female, spillover from female to male, facultative multi-sex (either group can maintain the pathogen), and obligate multi-sex (both groups are required to maintain the pathogen). The red dot represents the estimated value for *R*_0,*males*_, *R*_0,*females*_ (2.01, 0.42) for the WMHV system, using parameters from the best-fitting *6-group DD* model, predicting that males maintain WMHV in the system, with spillover occurring from males to females.

## Discussion

Identifying both the routes of disease transmission, and identifying the key individuals or groups that drive transmission in natural populations, is notoriously hard (25,26). One way to identify and quantify potential transmission routes is through the fitting of mechanistic models to epidemiological data, where we can estimate the magnitude of possible transmission-relevant parameters in the process (20). Here we built a series of epidemiological models of wood mouse herpesvirus (WMHV) transmission, each reflecting different hypotheses about the transmission processes within and between demographic groups in wild populations of the wood mouse, *Apodemus sylvaticus*, based on our previous results (15). We show that WMHV dynamics are dominated by transmission from males, particularly male-to-male transmission, and that it is likely that males can maintain the virus on their own, with females being little more than a ‘spillover’ group, playing a very small role in transmission in its natural host.

Despite being a standard model of herpesvirus infection biology and immunology in the laboratory, how WMHV and related viruses (e.g., MHV-68) transmit in the wild has been a long-standing mystery. It was generally believed that gammaherpesviruses transmit through close contact (13). However, previous research in laboratory studies in which infected and uninfected female mice were kept together in the same cage failed to result in transmission, suggesting that infection does not seem to happen through inhalation of virus particles suspended in the air. As such it has been suggested that the presence of males and their associated natural behaviours could be important in viral transmission, with possibly transmission occurring through sexual contact. For example, Francois *et al*. (2013) experimentally demonstrated herpesvirus transmission from infected females to naive males after sexual contact but not vice versa (17). However, an experiment with a different type of virus, Sin Nombre hantavirus, showed that natural transmission among mice in cages was uncommon and only occurred among males, but not females or between sexes (27).

Our results, which quantify natural transmission rates in the wild, confirm the importance of males for driving herpesvirus infection; the highest transmission rate related to males infecting other males, followed by male-to-female transmission. This dominance of males in driving transmission may arise from physiological (i.e. hormonal or immune-mediated) processes, behavioural differences, or both. The observed sex bias in our results and other empirical studies (15–17), may be due to infected males being more infectious than females. Suppression in immune responses are known to be a product of testosterone increases in reproductive males (28–30) and corticosteroids have been suggested to reduce the production of antibodies (31). In terms of behavioural explanations, previous studies on rodent viral infections have shown a higher prevalence in males for hantavirus (32) and cowpox (33), and it has been proposed that those differences are due to higher territorial aggression and travel distances in males. Thus, male dominance behaviours, such as scent-marking and biting, may constitute important transmission routes via urine and saliva, respectively. Laboratory experiments highlight the nose as an important point of viral entry (34), because among inoculation routes, intranasal is highly effective for infecting mice (35). Since WMHV prevalence in wood mice seems to be independent of breeding season (16) and previous studies have concluded that scent marking, at least in voles, could also convey identity and the frequency of scent marking was not always related to mate choice (36,37); scent-marking behaviour could be the main transmission route(15). As such we note that the relatively high rates of male to female transmission that we observe does not have to be due specifically to sexual contact but also scent-marking behaviour, since the latter is used to advertise current reproductive condition, attract mates, or merely individual identification (36,37). If scent-marking is the primary route of WMHV infection, then the relatively low contribution of females to transmission would be explained if females do not participate in this behaviour. However, few studies have considered scent marking among female wood mice. In *Mus musculus*, non-breeding females have been show to place fewer marks compared to breeding females (38), although female *Mesocricetus auratus* (39,40) and *M. musculus* (38) do use the frequency and placement of scent marks in competition with female conspecifics. Clarification of sex-specific scent marking behaviour in wild wood mice, along with assays of viral load in male and female urine, would help clarify the observed sex-specific differences in transmission.

To support the suggestion that transmission in these natural populations is not driven by sexual contacts, we found that a density-dependent transmission function best fit the observed seroprevalence patterns. This implies that frequency-dependent transmission, typically assumed for sexually transmitted pathogens (41), was not driving herpesvirus infection in this population. In addition, theory predicts that sexually transmitted infections would be female biased due to a higher variance in mating success among males (42), therefore exclusively sexual transmission is unlikely in polyandrous species, such as the wood mouse (43). However, previous research has emphasized that WMHV in wild wood mice shows no strong density-dependent patterns because prevalence was not significantly influenced by either the influx of juveniles during the breeding season or overall population density (16). Additionally, Knowles *et al*. (15) showed that population density increased from July onwards, but herpesvirus prevalence remained low. If prevalence is male biased (15,16) and infection is density-dependent, then transmission is likely to be related to the density of the high-risk group, rather than of the total population (44).

The data used in this study are an extension of that used by Knowles et al. (15) who found a marked, repeatable, seasonal variation in wood mice prevalence in one of the field sites (Haddon Wood) in 2009 and 2010, identifying an increase in prevalence between December and May. Our findings show that this pattern is also repeated in the two subsequent years, 2011 and 2012, in males and in 2012 in females. Thus, in accordance with Knowles *et al*., most of the transmission could occur during early spring previous to breeding season, in which male home range size increases (45), contact rates between reproductive males are higher compared to other times of the year and territories are established.

Regarding limitations of our study, antibody detection is not the best test for determining herpesvirus infection because maternal antibodies to the virus are inherited from mother to offspring and persist into young adulthood (15). Therefore, we excluded juveniles from our study; however, we acknowledge that using a PCR diagnostic method could refine our WMHV prevalence results. Furthermore, based on previous findings on murine gamma herpesvirus pathogenesis, our model did not assume disease-induced mortality due to infection (7,46). Nonetheless, previous work has found some evidence for a negative association between herpesvirus infection and recapture duration in adults, implying that infection may reduce life expectancy (15).

In summary, our study sheds light on the transmission mechanisms of a natural herpesvirus (WMHV) of wild wood mice by suggesting that transmission is density-dependent and mainly male-driven. Most of the virus transmission could therefore occur through scent-marking behaviour, before the breeding season, when males are establishing territories and home range sizes increase. Females play a less important role and could get infected predominantly by contact with scent-marked territories by males and, less likely, by other female conspecifics. Overall, we highlight the value of long-term longitudinal data, coupled with biologically meaningful mechanistic models, to elucidate key aspects of transmission biology, which otherwise would be hard to detect in the wild or the lab.

## Data accessibility

R code and data for reproducing the main results is available in the Dryad Digital Repository: https://datadryad.org/stash/share/wd9Lqi6F1nHOWJeGwoYov2iu7Ts4SYbEue-QjleuvFo

## Authors’ contributions

AB collected the field data, DE and AF designed the model, DE parameterized the model and drafted the manuscript, and DE, AF and KG analysed the results. All authors contributed to revisions.

## Acknowledgements

We would like to thank those involved in the fieldwork that generated the data used in this study, and the landowners for permission to carry out the work on their land.

